# Neurons in the Human Substantia Nigra Respond to Cognitive Boundaries and Predict Memory

**DOI:** 10.64898/2026.02.04.703556

**Authors:** Lin Shi, Anamar Flores, Lydia Shimelis, Yan Liu, Chenguan Jiang, Jianguo Zhang, Fangang Meng, Jie Zheng

**Author notes:** These authors contribute equally to the work (Lin Shi, Anamar Flores). Correspondence to: Jie Zheng,; Fangang Meng.

## Abstract

Segmenting mnemonic episodes from continuous experience is a key aspect of human episodic memory. The brain constantly forms predictions about what will happen next based on previous experience and knowledge, and prediction errors are thought to signal when a new event begins (cognitive boundaries). Dopamine has been closely linked to prediction error signals, yet it remains unknown how human midbrain neurons are modulated by cognitive boundaries and how their responses influence memory. To address these questions, we recorded activity of individual neurons in the human substantia nigra, a critical brain structure for dopamine production and regulation, while participants undergoing deep brain stimulation surgery watched a series of clips embedded with cognitive boundaries and performed a recognition memory task. We found that neural activity in the substantia nigra was robustly modulated by cognitive boundaries during clip viewing. Moreover, a subset of these boundary-responsive neurons also differentiated novel from familiar images during recognition, and their firing rates were indicative of participants’ memory success. These findings reveal that neurons in the human substantia nigra carry boundary- and novelty-related signals consistent with prediction error mechanisms that influence the encoding and retrieval of episodic memories.

## Introduction

Episodic memory is essential to how we understand the world and construct our identity, shaping experiences ranging from major life events to everyday routines. The brain captures individual moments and links them into detailed, coherent sequences that can later guide decisions, expectations, and future behavior. This ability to structure continuous experience into meaningful events is central to how memories are stored and retrieved. Transitions between these events, termed cognitive boundaries, mark points where ongoing experience is segmented into distinct units in memory. However, an unresolved question in the study of human memory is how the brain encodes ongoing experience at cognitive boundaries into representations that can be used effectively at a later time.

Event segmentation theory^1^ provides a framework for how the brain processes information and learns from experiences, and it also offers a theoretical model of how memorable moments (i.e., events) are segmented from continuous interaction with the world at cognitive boundaries. When incoming sensory information deviates from what is expected, a prediction error is generated. This error signal indicates that the original prediction was inaccurate or incomplete. In turn, prediction errors drive learning about new information or events by signaling the need to update existing predictions and memory representations^2–4^.

Dopamine is a neurotransmitter involved in signaling prediction errors. Dopaminergic neurons in the substantia nigra and ventral tegmental area release dopamine in response to prediction errors, thereby reinforcing or adjusting ongoing learning processes^5^. Dopaminergic neurons in the substantia nigra and ventral tegmental area have been extensively studied in the context of reward expectation errors^6–8^, odd-ball detection^9^, and decision making^10^. However, much less is known about their role in declarative memory beyond responses to novel stimuli in conditioning^11^ and recognition^12^ paradigms.

It has been proposed that the dopaminergic system and the hippocampus form a multisynaptic loop, in which the hippocampus sends novelty signals to dopaminergic neurons in the substantia nigra and ventral tegmental area. In turn, these midbrain neurons then send signals back to the hippocampus, where dopamine enhances synaptic plasticity via hippocampal dopamine receptors^13,14^. Previous work further shows that neurons in the human medial temporal lobe, including the hippocampus, detect cognitive boundaries and use them to structure episodic memories^15^. Taken together, this suggests that the hippocampus-substantia nigra circuit may also support cognitive boundary detection. We hypothesize that cognitive boundary signals computed in the hippocampus activate dopaminergic neurons in the substantia nigra, and that the resulting dopaminergic feedback enhances mnemonic representations in the hippocampus at event boundaries. While the original loop encompasses both the substantia nigra and the ventral tegmental area, here we focus specifically on the substantia nigra by leveraging clinical access to this region in patients undergoing deep brain stimulation (DBS) implantation surgery. Additionally, we do not limit our discussion to dopaminergic neurons in the substantia nigra, as γ-aminobutyric acid (GABA)ergic neurons in this region can inhibit dopaminergic neurons^16^ and therefore play a direct role in the circuit mechanisms addressed by our hypothesis.

To test this, we record single neuron activity from patients undergoing DBS surgery while they watched video clips containing cognitive boundaries and performed a recognition memory task in the operating room. We assess neuronal responses in the substantia nigra at cognitive boundaries during clip viewing and to novel images during the subsequent scene recognition test. Finally, we ask whether neural activity in the substantia nigra was predictive of memory formation success for the associated events.

## Results

### Task, behavior and neural recording

Forty patients (see demographics in **Supplementary Table 1**) undergoing implantation of a DBS device in the subthalamic nucleus for the treatment of Parkinson’s disease or dystonia performed a memory task in the operating room (**Figure 1a**). Single neuron activity was recorded while they performed a memory task (**Figure 1b**). The task consisted of two parts: encoding (**Figure 1c-e**) and scene recognition (**Figure 1f-h**). During encoding, participants watched a series of novel movie clips, each lasting approximately 6 seconds (see *Methods*). Each clip was either a single continuous shot (**Figure 1e**, “virtual boundary clips”) or two different shots separated by a cognitive boundary (**Figure 1d**, “cognitive boundary clips”). Cognitive boundaries were defined as moments when the clip transitioned to an unexpected new scene that was contextually unrelated to the preceding content. For example, in the representative clip (**Figure 1d**), the cognitive boundary refers to the cut from checking the compass to various activities in an amusement park. As a comparison condition, we defined “virtual boundaries” as the time point 3 seconds after the onsets of virtual boundary clips, chosen to approximate the midpoint of the 6 second clips and to provide a time-matched reference point relative to the cognitive boundary condition. At the end of each clip, participants reported whether most of the scene took place “indoors” or “outdoors”, ensuring their engagement and providing a measure of encoding success. After watching twenty clips in a session, participants were instructed to complete a scene recognition memory test (**Figure 1f**). During scene recognition, participants were presented with single static frames and were instructed to indicate via button press whether each frame was “old” (from a watched clip) or “new” (not previously seen). Most participants completed two sessions of encoding and scene recognition tests, each with different clips. Of the 40 patients, 11 completed three sessions.

**Figure 1.**
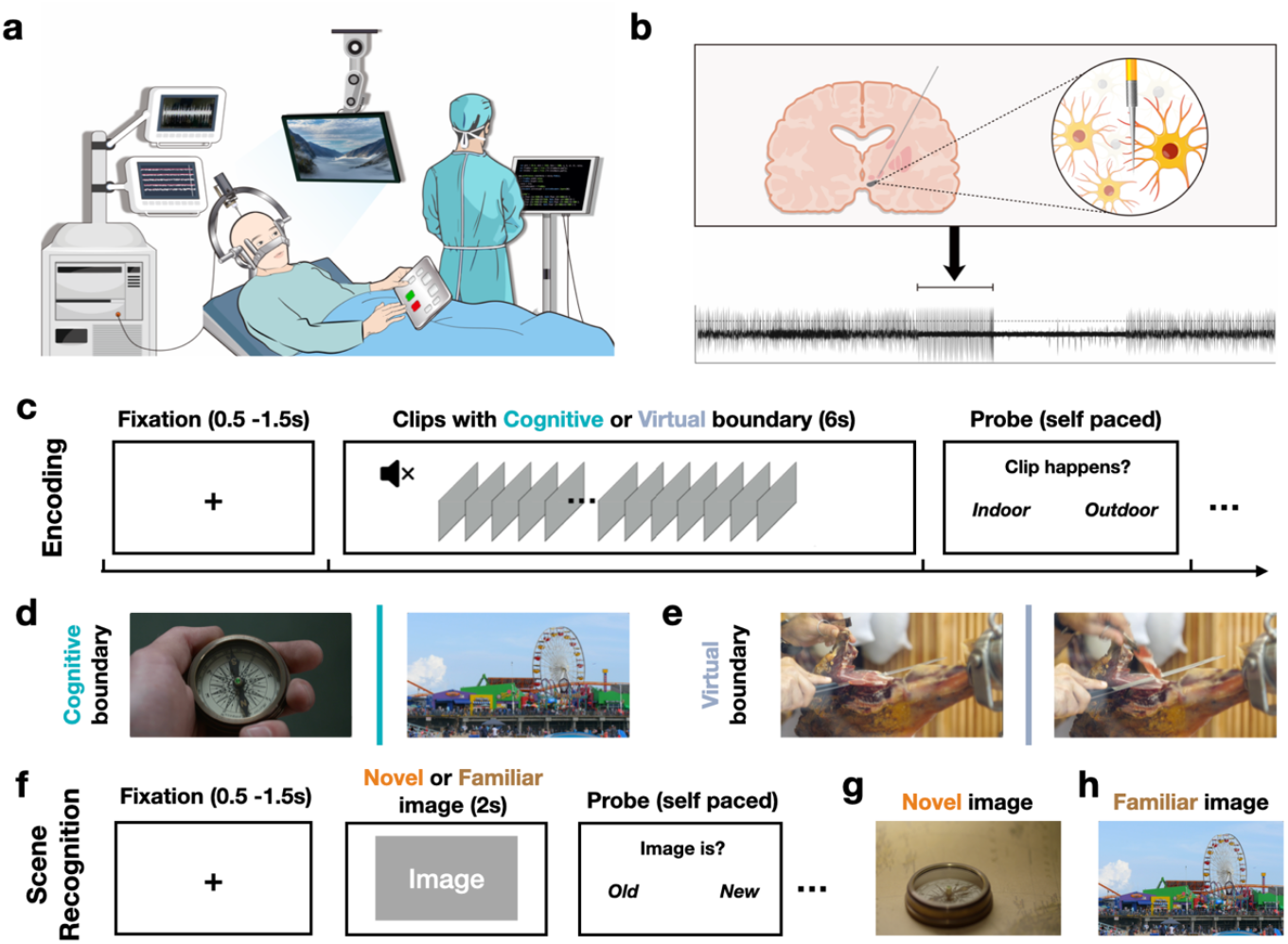
Experiment and task setup. **(a)** Schematic of the intraoperative experimental setup. **(b)** Illustration of microelectrode single unit recording in the substantia nigra: electrode placement and neurons near the electrode tip (top), and an example raw recording trace (bottom). **(c-h)** Schematic of the two-stage task. **(c)** Encoding. Participants viewed a series of silent video clips and, after each clip, indicated whether most of the clip took place indoors or outdoors. Clips either contained a cut between scenes from different movies (cognitive boundary clips) or contained no cut (virtual boundary clips). For virtual boundary clips, a “virtual boundary” time point was defined at 3 seconds after clip onset (the clip midpoint) and used as the alignment point for analyses comparing virtual and cognitive boundaries. **(d**,**e)** Example frames spanning a cognitive boundary (cut) and a virtual boundary (no cut). **(f)** Scene recognition. Participants viewed a static image and reported whether it was “old” (previously shown during encoding) or “new.” (**g**,**h**) Examples of novel and familiar images. Task prompts are shown here in English; the original task instructions were presented in Chinese (see **Supplementary Figure 1**).

Participants performed well on the memory task in the operating room. During encoding, participants correctly answered the “indoor” or “outdoor” questions on 84.7% ± 9.6% (mean ± standard deviation, reported throughout) of trials, significantly above the chance level of 50% (*t*_*39*_ = 22.84, *p* = 0, one sample *t*-test with respect to chance level; **Figure 2a** dark grey dots). During scene recognition, participants correctly recognized familiar images or correctly identified novel images on 73.6% ± 10.8% of trials, again significantly above the chance level of 50% (*t*_*39*_ =13.81, *p* = 1.11×10^-16^, one sample *t*-test with respect to chance level; **Figure 2a** light gray dots). Participants also had higher accuracy (cognitive boundary clips: 82.5 ± 13.2%, virtual boundary clips: 64.7 ± 10.4%, *t*_*39*_ = 11.41, *p* = 5.37×10^-14^, two-tailed paired *t*-test; **Figure 2b**) but took a longer time (cognitive boundary clips: 1.35 ± 0.88 seconds, virtual boundary clips: 1.18 ± 0.72 seconds, *t*_*39*_ = 3.02, *p* = 0.004, two-tailed paired *t*-test; **Figure 2c**) to recognize images extracted from cognitive boundary clips compared to the ones extracted from virtual boundary clips.

**Figure 2.**
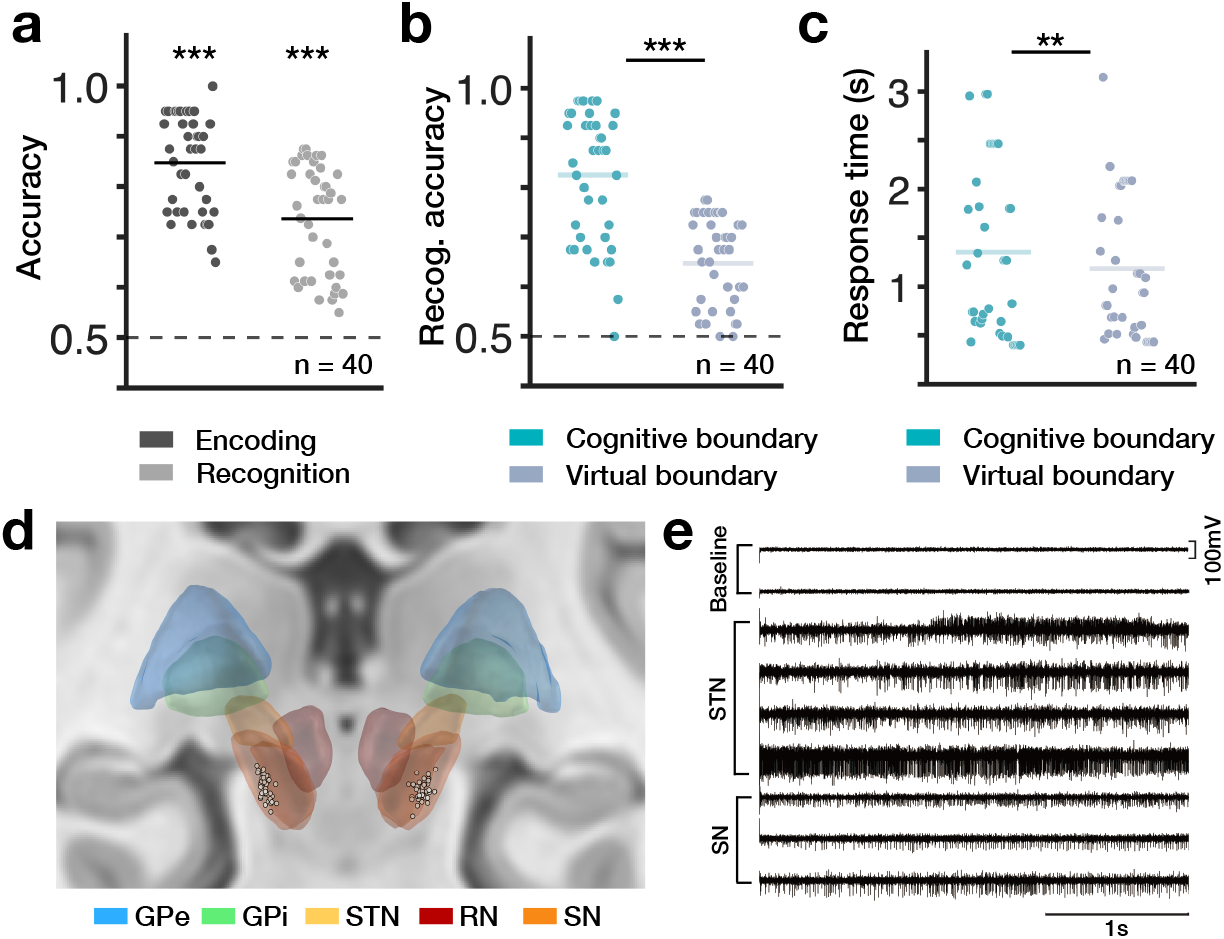
Behavioral results and recording locations. **(a)** Task performance for patients undergoing DBS implantation. Encoding accuracy (dark gray) is the proportion of trials in which participants correctly answered the indoor/outdoor question during encoding. Recognition accuracy (light gray) is the proportion of trials in which participants correctly judged images as old/new during recognition. **(b)** Recognition accuracy for clips containing cognitive boundaries (blue) versus virtual boundaries (gray).**(c)** Response times for cognitive boundary (blue) versus virtual boundary (gray) clips. **(d)** Recording sites across 40 participants (participant demographics in **Supplementary Table 1**). Microelectrode locations are shown in MNI space on a template brain (see *Methods*) and plotted as individual dots, color-coded by region: blue, GPe (globus pallidus externus); green, GPi (globus pallidus internus); yellow, STN (subthalamic nucleus); red, RN (red nucleus); orange, SN (substantia nigra; including substantia nigra pars compacta and substantia nigra pars reticulata). MNI coordinates for all recording sites are provided in **Supplementary Tables 2** and **3. (e)** Example microelectrode recordings along the implantation trajectory.

Single-neuron activity was recorded from the substantia nigra while participants performed the memory task. The substantia nigra, located just below the subthalamic nucleus, is a common entry point for DBS implantation targeting the subthalamic nucleus. Its distinct electrophysiological properties (**Figure 2e**), with more regular and slower firing compared to the subthalamic nucleus, help define the subthalamic nucleus border during surgery (see *Methods* for details on region identification). We leveraged this opportunity to record neural activity when the probe was positioned in the substantia nigra. We identified 134 putative single neurons in total (3.3 ± 0.8 clusters per recording channel). The locations of recording sites (see Montreal Neurological Institute coordinates listed in **Supplementary Tables 2 and 3**) with putative neurons identified were plotted in the template brain (**Figure 2d**; see *Methods*). Neurons were well isolated as assessed quantitatively using spike sorting quality metrics (**Supplementary Figure 2**). Throughout the manuscript, we use the terms neuron, unit, cell interchangeably to refer to the putative single neuron recorded for this task.

### Neurons in the substantia nigra respond to cognitive boundaries

We first asked whether substantia nigra neurons modulate their firing rates at cognitive boundaries during clip watching. Among all 134 recorded substantia nigra neurons, 36 of them (36.6%, *p* = 0.008, compared to chance level using permutation test) showed significant changes in activity within the 1 second window following cognitive boundaries relative to the 1 second baseline period preceding them (see *Methods*). We refer to these neurons as *boundary cells*. Boundary cells with increased post-boundary firing rates were classified as *type-I boundary cells* (see example in **Figure 3a**), whereas those with decreased firing were classified as *type-II boundary cells* (see example in **Figure 3b**). Among all the substantia nigra neurons recorded, we found 22 of them (16.4%, *p* = 0.012, compared to the chance level, permutation test) as type-I boundary cells (10.4%, *p* = 0.016, compared to the chance level, permutation test) and 14 of them as type-II boundary cells.

**Figure 3.**
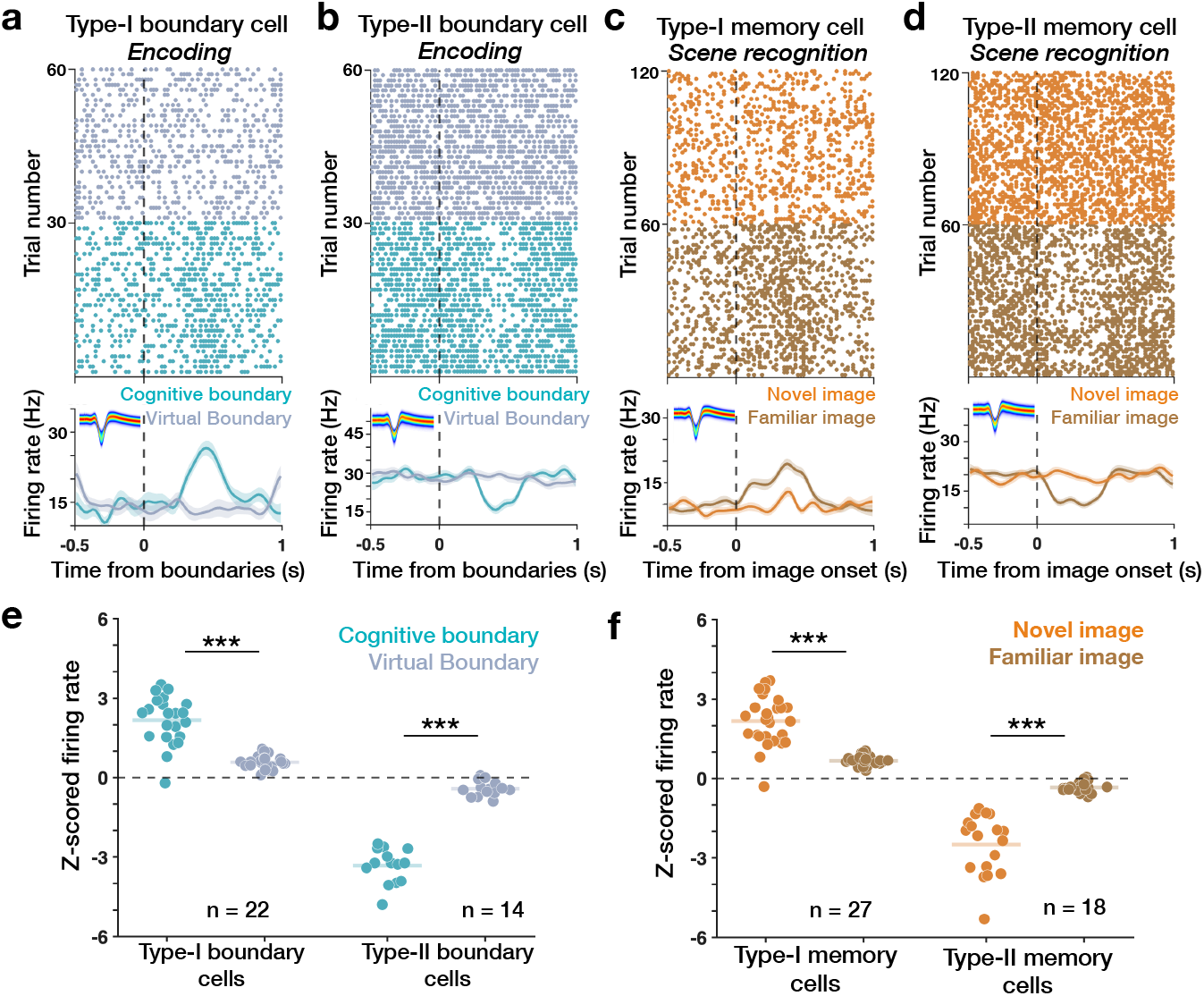
Boundary and memory cell types in the substantia nigra. **(a, b)** Example boundary cells.**(a)**Raster plot (top) and peristimulus time histogram (bottom) from a representative type-I boundary cell recorded in the substantia nigra, showing elevated firing following cognitive boundaries (blue) relative to virtual boundaries (gray). **(b)** Same as **(a)**, for a representative type-II boundary cell, showing suppressed firing following cognitive boundaries (blue) relative to virtual boundaries (gray). **(c, d)** Example memory (novelty) cells. **(c)** Raster plot (top) and peristimulus time histogram (bottom) from a representative type-I memory cell, showing elevated firing following novel image onsets (orange) relative to familiar image onsets (brown). **(d)** Same as **(c)**, for a representative type-II memory cell, showing suppressed firing following novel (orange) relative to familiar (brown) image onsets. **(e)** Population responses: baseline-normalized (z-scored) firing rates following cognitive boundaries (blue) and virtual boundaries (gray) are shown for all type-I boundary cells (n = 22) and type-II boundary cells (n = 14). **(f)** Population responses: baseline-normalized (z-scored) firing rates following novel (orange) and familiar (brown) image onsets are shown for all type-I memory cells (n = 27) and type-II memory cells (n = 18).

Boundary-related responses were selective to cognitive boundaries rather than generic properties of clip structure. For each boundary cell, we computed the z-scored firing rate in the 1 second window following cognitive boundaries by normalizing it to firing rates in the 1 second pre-boundary baseline. We repeated this procedure for clips without cognitive boundaries, aligning neural activity to virtual boundaries positioned 3 seconds after clip onsets. Comparing neural responses to cognitive boundaries versus virtual boundaries revealed that type-I boundary cells reliably exhibited stronger firing-rate increases to cognitive boundaries (z-scored firing rates: 2.17 ± 0.91) than to virtual boundaries (z-scored firing rates: 0.58 ± 0.25; *t*_21_ = 7.80, *p* = 1.25×10^-7^; **Figure 3e**). Conversely, type-II boundary cells showed larger firing rate decreases following cognitive boundaries (z-scored firing rates: -3.33 ± 0.66) relative to virtual boundaries (z-scored firing rates: -0.41 ± 0.23; *t*_13_ = -16.58, *p* = 3.99×10^-10^). Together, these results reveal that a discrete subset of substantia nigra neurons encodes cognitive boundaries via brief, boundary-locked modulations in firing rate.

### Neurons in the substantia nigra differentiate novel versus familiar images

We next examined how substantia nigra neurons responded during the scene recognition task, in which participants identified previously viewed images as “old” and unseen images as “new.” For each neuron recorded in the substantia nigra, we compared its average firing rate within the 1 second time window after the onset of a novel image and compared it with its firing rate within the 1 second time window before the image appeared. This analysis revealed that 45 of 134 neurons (33.6%, *p* = 0.002, permutation test) showed significant changes in activity when novel images were presented, classified as *memory cells*. We refer to neurons that increased their firing during novel images presentation as *type-I memory cells* (see example in **Figure 3c**) and those that decreased their firing as *type-II memory cells* (see example in **Figure 3d**). We observed a slightly higher ratio of type-I memory cells than type-II memory cells. Among all the recorded substantia nigra neurons, 27 of them (20.1%, *p* = 0.009, compared to the chance level, permutation test) were classified as type-I memory cells,while 18 of them (13.4%, *p* = 0.011, compared to the chance level, permutation test) as type-II memory cells.

We then asked whether these 45 memory cells also responded to familiar images. We found that only a small number did. Of the cells recorded, 2 of 27 type-I memory cells (7.4%) and 1 of 18 type-II memory cells (5.6%) showed significant firing rate changes following the onsets of familiar images. To compare responses more directly, for each memory cell, we computed the average firing rate within 1 second time window after both novel and familiar images, and z-scored against their firing rates within the 1 second pre-image period as the baseline. We found that type-I memory cells consistently showed stronger firing rate increases to novel images (z-scored firing rate: 2.17 ± 0.93) than to familiar ones (z-scored firing rate: 0.68 ± 0.17; novel images versus familiar images: *t*_26_ = 8.81, *p* = 2.75×10^-9^; **Figure 3f**). The same pattern also held for type-II memory cells, which showed larger firing rate decreases to novel images (z-scored firing rate: -2.49 ± 1.13) than to familiar images (z-scored firing rate: -0.33 ± 0.20; novel images versus familiar images: *t*_17_ = -7.97, *p* = 3.87×10^-7^).

### Substantia nigra neurons respond to both cognitive boundaries and novel images

Given that cognitive boundaries are a type of novelty signal, we then asked whether neurons that respond to novel images during scene recognition also respond to cognitive boundaries during encoding. We found that boundary cells and memory cells were highly overlapped, with 26 of 134 recorded substantia nigra neurons (19.4%, *p* = 0.003, compared to null distribution, permutation test) responsive to both cognitive boundaries during encoding and novel images during scene recognition.

Within this overlap group, 16 cells increased firing following both cognitive boundaries and novel images (*type-I overlap*; **Figure 4a**) and 10 cells decreased firing to both events (*type-II overlap*; **Figure 4b**). Relative to their respective parent populations, type-I overlap cells comprised 16 of 33 type-I cells (48.5%) and type-II overlap cells comprised 10 of 22 type-II cells (45.4%). Note that there were no overlaps between type-I boundary cells and type-II memory cells, nor type-II boundary cells and type-I memory cells. For both type-I overlap cells and type-II overlap cells, the firing rate changes following cognitive boundaries during encoding and novel images during scene recognition were highly correlated (**Figure 4c**: type-I overlap cell, *r* = 0.77, *p* = 4.39×10^-4^, Pearson’s correlation; **Figure 4d**: type-II overlap cell, *r* = 0.80, *p* = 5.32×10^-3^, Pearson’s correlation).

**Figure 4.**
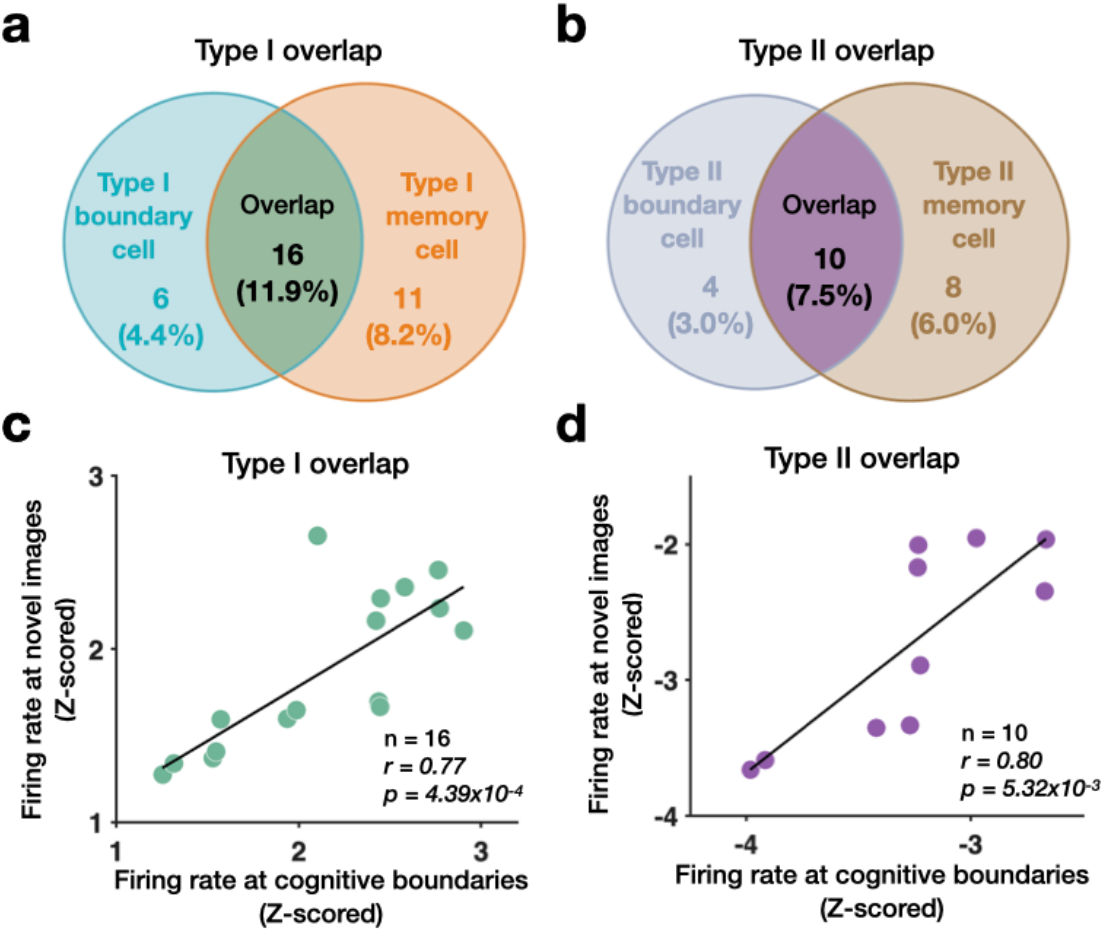
Relationship between boundary cells and memory cells. **(a, b)** Overlap between boundary cells and memory cells for type-I **(a)** and type-II **(b)** populations. Numbers indicate cell counts in each category (boundary-only, memory-only, and overlap), and percentages indicate each category as a fraction of all substantia nigra neurons recorded. (**c, d**) For overlap neurons, relationship between boundary-evoked and novelty-evoked activity. Scatter plots show, for each neuron, mean baseline-normalized (z-scored) firing rates averaged in a 1 second window following cognitive boundaries (x-axis) versus a 1 second window following novel image onsets (y-axis) for type-I overlap cells **(c)** and type-II overlap cells **(d)**.

We then asked how type-I overlap cells and type-II overlap cells differ from each other. First, we examined their anatomical distribution by plotting the recording locations of type-I (green spheres) and type-II (purple spheres) overlap cells on a template brain in MNI space (ICBM 2009b nonlinear asymmetric MNI template^17^ and DBS Tractography Atlas; Middlebrooks, 2020^18^), using the coordinates in **Supplementary Figure 1**, with substantia nigra pars compacta (SNc) plotted in dark gray and substantia nigra pars reticulata (SNr) in light gray (**Figure 5a**). Note that type-I overlap cells and type-II overlap cells recorded on the left hemisphere have been projected to the right hemisphere by taking the absolute value of their MNI coordinates in the lateral-medial axis. We found that type-I overlap cells tend to locate in more ventral (type-I: -15.09 ± 1.13; type-II: -13.42 ± 0.94; type-I versus type-II: *F*(1,24) = 15.19, *p* = 6.83×10^-4^, one-way ANOVA), medial (type-I: 9.29 ± 0.49; type-II: 11.06 ± 0.49; type-I versus type-II: *F*(1,24) = 80.53, *p* = 3.90×10^-9^, one-way ANOVA), and posterior (type-I: -18.52 ± 0.94; type-II: -15.69 ± 1.74; type-I versus type-II: *F*(1,24) = 29.33, *p* = 1.46×10^-5^, one-way ANOVA) subregions within the substantia nigra compared to type-II overlap cells (**Figure 5a**). Similar anatomical distribution is also observed when extending the analyses to all the type-I boundary cells versus type-II boundary cells, and type-I memory cells versus type-II memory cells (**Supplementary Figure 3**). Relative to type-II boundary cells, type-I boundary cells were located more ventrally (type-I boundary: -15.03 ± 1.07; type-II boundary: -13.78 ± 1.04; *F*(1,34) = 13.33, *p* = 8.69×10^-4^), more medially (type-I boundary: -9.40 ± 0.65; type-II boundary:

**Figure 5.**
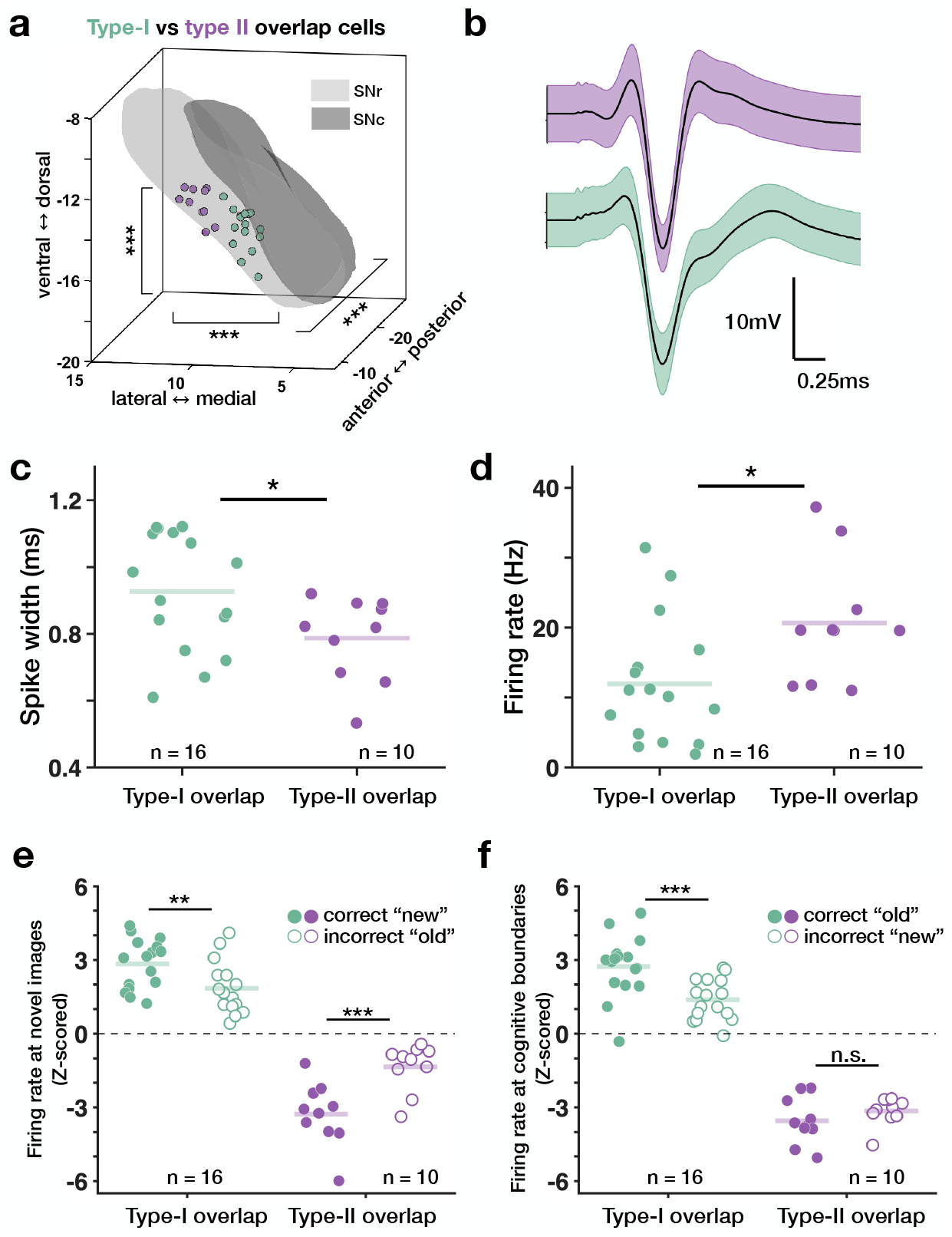
Comparison of type-I and type-II overlap cells. **(a)** Microelectrode recording locations of overlap cells plotted in MNI space and color-coded by subtype (type-I, green; type-II, purple). To pool hemispheres, left-hemisphere sites are mirrored to the right by taking the absolute value of the lateral (X) coordinate. SNc and SNr are shown in dark gray and light gray, respectively. **(b)** Mean spike waveform (± s.d.) for representative type-I (green) and type-II (purple) overlap cells. **(c)** Spike width, defined as the time between the two positive peaks of the waveform, for type-I and type-II overlap cells. **(d)** Mean firing rate across the entire recording session for type-I and type-II overlap cells. **(e)** Baseline-normalized (z-scored) firing rates in a 1-second window following novel image onset, shown separately for trials in which novel images were correctly judged “new” versus misclassified as “old,” for type-I and type-II overlap cells. **(f)** Baseline-normalized (z-scored) firing rates in a 1-second window following cognitive boundaries, shown separately for trials in which familiar images were correctly judged “old” images versus misclassified as “new,” for type-I cells and type-II overlap cells.

10.87 ± 0.52; *F*(1,34) = 50.79, *p* = 3.09×10^-8^), and more posteriorly (type-I boundary: - 18.45 ± 1.65; type-II boundary: -15.97 ± 1.56; *F*(1,34) = 20.12, *p* = 7.89×10^-5^) within the substantia nigra. Similarly, type-I memory cells were also located more ventrally (type-I memory: -14.98 ± 1.13; type-II memory: -13.75 ± 0.94; *F*(1,43) = 14.75, *p* = 3.99×10^-4^), more medially (type-I memory: 9.46 ± 0.61; type-II memory: 10.86 ± 0.46; *F*(1,43) = 67.70, *p* = 2.23×10^-10^), and more posteriorly (type-I memory: -18.47 ± 1.52; type-II memory: -16.01 ± 1.41; *F*(1,43) = 30.07, *p* = 2.05×10^-6^) than type-II memory cells.

The substantia nigra contains two main neuronal populations: inhibitory GABAergic neurons, which are predominantly located in the SNr, and dopaminergic neurons in the SNc, which project broadly to regions such as the striatum, amygdala, and hippocampus^19–21^. Given this distinct anatomical and functional division, we asked whether type-I and type-II overlap cells might correspond to different underlying cell types. In general, in the substantia nigra, dopaminergic neurons have wider waveforms and lower firing rates compared to GABAergic neurons. Therefore, we assessed the waveform length by computing the time that elapsed between the two positive peaks of the waveform^8^. We found that type-I overlap cells (see example in **Figure 5b** green color; 0.93 ± 0.18 milliseconds) were on average characterized by longer waveforms (i.e. larger peak-to-peak width of spike waveform) compared to type-II overlap neurons (see example in **Figure 5b** purple color; 0.79 ± 0.13 milliseconds; type-I versus type-II: *F*(1, 24) = 4.81, *p* = 0.038, one-way ANOVA; **Figure 5c**). In addition, type-I overlap cells had lower mean firing rates (averaged across the entire recording session) compared to type-II overlap cells (type-I overlap neurons: 11.91 ± 8.85 Hz; type-II overlap neurons: 20.64 ± 8.88 Hz; type-I versus type-II: *F*(1, 24) = 5.97, *p* = 0.022, one-way ANOVA; **Figure 5d**). Additionally, 11 of 16 (68.8%) type-I overlap cells satisfied the criteria for dopaminergic neurons reported in previous studies^8,12,22,23^, with its peak-to-peak width of spike waveform longer than 0.8 milliseconds and mean firing rates lower than 15Hz.

### Neurons responsive to both cognitive boundaries and novel images predict participants’ memory performance

We next examined whether the neural responses of type-I and type-II overlap cells to cognitive boundaries and novel images were related to participants’ memory performance. We began with the scene-recognition task. For each overlap cell, we calculated the firing rate in the 1 second window following novel images, z-score normalized to the 1 second baseline window before the image, and compared activity between trials in which novel images were correctly labeled as “new” and those in which they were mistakenly judged as “old.” Type-I overlap cells showed stronger increases in firing relative to baseline when participants correctly identified novel images (2.84 ± 1.01) than when they misclassified them “old” (1.86 ± 1.05; correct versus incorrect: *t*_15_ = 3.48, *p* = 0.0034, two-tailed paired *t*-test; **Figure 5e** green color). Conversely, type-II overlap cells decreased their firing rate more strongly when participants accurately reported novel images as “new” (−3.28 ± 1.28) versus mistakenly as “old” (−1.35 ± 0.95; correct versus incorrect: *t*_9_ = -4.90, *p* = 8.52×10^-4^, two-tailed paired *t*-test; **Figure 5e** purple color).

We then asked whether overlap cell activity at cognitive boundaries during encoding predicted later recognition performance. Using the same approach as above, we computed z-scored firing rates in the 1 second window after each cognitive boundary and compared activity between trials where participants later correctly recognized the tested images and those where they did not, separately for novel and familiar images. For type-I overlap cells, firing rate changes following cognitive boundaries were stronger for familiar images that were later correctly recognized as “old” (2.73 ± 1.25) compared to the those were forgotten (1.38 ± 0.83; correct versus incorrect: *t*_15_ = 5.14, *p* = 1.22×10^-4^, two-tailed paired *t*-test; **Figure 5f** green color). In contrast, type-II overlap cells had no difference in firing changes between correctly recognized familiar images (−3.55 ± 0.95) and forgotten ones (−3.15 ± 0.57; correct versus incorrect: *t*_9_ = -1.50, *p* = 0.17, two-tailed paired *t*-test; **Figure 5f** purple color).

## Discussion

In this study, participants undergoing DBS implantation surgery performed a memory task involving dynamic video clips while single neuron activity was recorded in the substantia nigra. During encoding, when participants viewed continuous video clips, a subset of substantia nigra neurons showed increased (type-I boundary cells) or decreased (type-II boundary cells) firing rates at cognitive boundaries (i.e., visual cuts involving scene transitions within the clip) compared to the baseline period right before the cognitive boundaries. During memory retrieval, when participants were instructed to identify tested frames as “old” or “new”, a subset of substantia nigra neurons exhibiting increased (type-I memory cells) or decreased (type-II memory cells) firing rates to novel images relative to the fixation period. Boundary cells and memory cells were substantially overlapped, and the firing rates of the overlapping cell population following cognitive boundaries during encoding and during image presentation in the scene recognition were highly correlated and predictive of participants’ memory outcomes.

Memory has been studied most extensively in cortical regions, particularly the medial temporal lobe. How do our findings in the substantia nigra relate to the existing medial temporal lobe dependent memory system? Here we report that neurons in the human substantia nigra change their firing rates (type-I memory cells and type-II memory cells) when participants are presented with novel images and are asked to identify them as “new”. In particular, the type-I memory cells we identify in the substantia nigra behave similarly to the memory-selective neurons previously reported in the medial temporal lobe^24^, with both populations increasing their firing rates to novel relative to familiar images. Notably, type-I memory cells in the substantia nigra responded slower (∼300ms) to the novel images compared to the memory cells reported in the medial temporal lobe (∼250ms). This latency difference is compatible with the Lisman and Grace model^13,14^, in which a hippocampal novelty signal transiently excites dopaminergic neurons in the substantia nigra or ventral tegmental area. However, such comparison needs to be further tested, as the latency analyses were performed on two different patient populations (DBS patients versus epilepsy patients) with different recording systems (Neuro Omega versus Neuralynx ATLAS).

What functional roles may substantia nigra neurons play in supporting cognitive boundary detection? Classic work has established that midbrain dopamine neurons encode reward prediction errors, or discrepancies between expected and received outcomes^25^. Consistent with this, recordings in humans show that substantia nigra neurons respond to unexpected financial rewards^8^ and to deviant stimuli in odd ball paradigms^9^, both of which are forms of prediction error. Theories of event perception and memory propose that the brain uses such prediction errors to segment continuous experience, with cognitive boundaries occurring when hippocampal predictions about ongoing events are violated by incoming sensory input^26,27^. Within this framework, our observation that neurons in the human substantia nigra change their firing rates following cognitive boundaries (type-I boundary cells and type-II boundary cells, **Figure 3**), similar to boundary cells previously identified in the medial temporal lobe^15^, suggests that boundary-related responses in the substantia nigra may reflect a prediction error signal downstream of hippocampal boundary responses, consistent with models proposing that high prediction error at event boundaries reorganizes memory representations. This interpretation is further supported by the temporally delayed boundary responses in the substantia nigra compared to boundary responses in the hippocampus previously mentioned. As with the novelty signal, however, this timing difference needs to be validated within the same patient population in the future.

Why do neurons in the substantia nigra respond to both cognitive boundaries during encoding and novel images during scene recognition? Both events correspond to violations of expectation. Cognitive boundaries occur when incoming sensory information violates participants’ predictions about the ongoing scene, and novel images are items that have not been previously encountered. In line with this, the overlap between memory cells and boundary cells is highly structured, as neurons that increase firing at cognitive boundaries also increase firing to novel images (type-I overlap cells, **Figure 4a** and **4c**), whereas neurons that decrease firing at cognitive boundaries also decrease firing to novel images (type-II overlap cells, **Figure 4b** and **4d**). We observe no cross-polarity overlap between novelty and boundary coding, (i.e., no overlaps between type-I boundary cells and type-II memory cells, no overlaps between type-II boundary cells and type-I memory cells). This pattern is consistent with the idea that substantia nigra neurons encode a common novelty-or prediction-error-related response that generalizes across distinct task phases and stimulus types. In contrast, in the hippocampus, novelty and boundary responses are largely segregated into distinct cell groups with minimal overlap^15^. This pattern suggests that the substantia nigra may act as a centralized processor that integrates multiple forms of hippocampal novelty-related signals, while hippocampal regions maintain more differentiated, content-specific representation of event structure.

Why do we observe both type-I and type-II overlap cells in the substantia nigra? These two populations differ in several respects. First, type-I overlap cells exhibit broader spike waveforms than type-II overlap cells (**Figure 5c**). Second, type-I overlap cells have a lower mean firing rate than type-II overlap cells (**Figure 5d**). In addition, their firing rate changes show opposite polarities in response to cognitive boundaries and novel images, with these shifts differing systematically between the two cell types (**Figure 3e** and **3f**). Third, type-I and type-II overlap cells also appear at different anatomical subdivisions, with type-I overlap cells largely located in the ventral, medial and posterior part of the substantia nigra compared to the type-II overlap cells (**Figure 5a**). Taken together, the differences in spike width, mean firing rate, and anatomical distribution suggest that type-I overlap cells correspond primarily to dopaminergic neurons in the substantia nigra pars compacta, while type-II overlap cells might represent putative GABAergic neurons in the substantia nigra pars reticulata. The absence of type-II overlap cells (i.e., more GABAergic like neurons) in prior hippocampal recordings may reflect the relatively low proportion of GABAergic neurons in the hippocampus^28,29^ compared to substantia nigra pars reticulata^30^.

Why do boundary and novelty responses vary as a function of participants’ memory performance? During scene recognition, firing rates of type-I overlap cells and type-II overlap cells differed in response to novel images when participants correctly identified as “new” versus those misclassified as “old” (**Figure 5e**). However, during encoding, only the boundary-evoked activity in type-I overlap cells predicted later recognition of familiar images whereas type-II overlap cells did not differentiate later memory outcomes (**Figure 5f**). These results support a functional dissociation. Theories of the hippocampal-midbrain loop propose that dopaminergic modulation of hippocampal circuits enhances synaptic plasticity for behaviorally significant events, such as those that are rewarding, motivationally relevant, or that capture attention^13,14^. Indeed, type-I overlap cells/dopaminergic-like signals provide an event-level, boundary-locked prioritization that promotes plasticity across entire episodes and also scales with successful assessment of item novelty. In the context of the hippocampal-midbrain loop, these responses may trigger plasticity in hippocampal circuits and additional dopaminergic release, strengthening memory representations for novel stimuli and ultimately supporting the predictive relationship between boundary-related activity during encoding and later recognition of familiar images. However, type-II overlap cells/GABAergic-like signals likely reflect an inhibitory counterpart that decreases with unexpected input and relates to accurate novelty judgments but does not, on its own, predict subsequent episodic retention. A critical next step is to modulate these two groups of cells and causally test their influence on boundary detection and subsequent memory recall.

## Methods

### Participants

Between December 2019 and December 2021, 75 Chinese patients volunteered for this study and provided their informed consents. The inclusion criteria were as follows: 1) aged between 50 and 75 years; 2) diagnosed with idiopathic PD; 3) indicated for bilateral Subthalamic Nucleus (STN) DBS surgery; 4) agreement to participate in the intraoperative task and provision of written informed consent signed by the patient or their legal guardian. The participants performed the task in the operating room prior to the implantation of the DBS device. Participants whose preoperative Mini-mental State Examination (MMSE) score was less than 22, whose Montreal Cognitive Assessment (MoCA) score was less than 18, who could not cooperate with the experimenters either in the training or in the real surgery, or whose intraoperative electrophysiological signal quality was so poor that no stable spike discharges were recorded were excluded. Therefore, 40 participants were included in the study (12 females, mean age = 57.10 ± 10.11 years, see participants’ demographics in **Supplementary Table 1**). The study protocol was approved by the Institutional Review Board at Beijing Tiantan Hospital (No. KY2019-097-02). The location of the implanted electrodes was solely determined by clinical needs.

### Electrophysiological recordings

All patients underwent a bilateral, frame-based STN DBS surgery. Stereotactic planning was carried out with preoperative 3.0T MRI merged with stereotactic CT scan obtained on the day of surgery using the SurgiPlan software (Elekta, Sweden). Coordinates of the DBS targets and the entry points were calculated on the software using the direct targeting method as described elsewhere^31^. Local infiltration anesthesia was conducted (1% lidocaine together with 0.5% ropivacaine infused surrounding the entry point on the frontal part).

The surgical procedures were described in previous reports^32^. Briefly, after anesthesia and disinfection, incisions were made on the scalp near the entry point of DBS trajectories. A 1.5cm burr hole was made on each side of the skull. Microelectrode recording was conducted with a single tungsten microelectrode (400-700 kOhm, Neuroprobe, Alpha Omega, Israel) driven through the central passage on the electrode holder (Bengun, Alpha Omega, Israel) into the target area by a microdrive (Drive Headstage, Alpha Omega, Israel) with a step size of 100-200 μm. The dura puncture technique was employed to lessen the cerebrospinal fluid loss^33^. STN signals were recorded (Neuro Omega, Alpha Omega, Israel; see *Localization of substantia nigra for behavioral tasks during the DBS surgery*)^34^. The intraoperative tasks were performed when the neuronal firing signal of substantia nigra was stable and continuously recorded. After the task, the DBS electrodes (Model 3389/3389S, Medtronic, US; L 301, Pins, China) were implanted, with the lowest contact of the electrode being placed right on the boundary between the STN and substantia nigra.

### Localization of microelectrode recording sites

The microelectrode recording sites when the intraoperative behavioral tasks were performed were determined using the following methods.

#### Localization of substantia nigra for behavioral tasks during the DBS surgery

During STN-DBS surgery, microelectrode recording (MER) was employed to delineate the electrophysiological signatures of the STN and adjacent structures, including the substantia nigra (SN), as described elsewhere^35^. The STN was identified by its characteristic high-frequency, irregular neuronal discharges with background noise. As the microelectrode descended further, the transition from the STN to the SN was marked by a distinct shift in electrophysiological activity: the SN exhibits lower-frequency (2–6 Hz), regular tonic firing patterns and a reduction in background noise. To further corroborate SN localization, high-frequency and low-intensity microstimulation (0.1 mA, 200 Hz, 60 us) was applied at the microelectrode tip. Transient suppression of neuronal firing in the SN following stimulation served as an additional functional marker (**Figure 1h**). Final electrophysiological recording sites for behavioral testing were conducted at depths where stable, stereotyped nigral unit activity was observed.

#### Postoperative Localization of MER Sites

The post-operative reconstruction of the DBS electrode was performed using the Lead-DBS toolbox (v2.6) as described previously^18^. Following DBS implantation, high-resolution CT scans (0.625 mm slice thickness) were acquired without gap to visualize the electrode trajectories. These CT images were co-registered with preoperative MRI (T1/T2-weighted sequences) using rigid and affine transformations in subject-specific space. To standardize anatomical localization, individual brain structures were normalized to a common stereotactic space (i.e., MNI space) using advanced linear registration tools. The fused imaging data enabled 3D reconstruction of the DBS electrodes relative to deep brain nuclei (STN and SN). By integrating intraoperative microelectrode recording depths with the normalized imaging coordinates (adding the depth difference obtained during the surgery relative to the tip of the first contact of the DBS electrodes in the DBS trajectory), the precise spatial locations of electrophysiological signal acquisition during behavioral tasks were determined. Finally, the recording sites were projected onto MNI space to derive standardized coordinates, facilitating group-level analysis and comparisons.

### Spike sorting and quality metrics of single units

Spike sorting was performed offline using a semi-automated template matching algorithm Osort^36^. See Zheng et al. 2022^15^ for more details on spike sorting procedures. We identified 146 neurons in this dataset across all brain areas considered. We excluded 12 neurons with low firing rates (< 0.5Hz) and analyzed the remaining 134 neurons. The quality of our spike sorting results was evaluated using our standard set of spike sorting quality metrics^37,38^ for all considered 134 putative single neurons (**Supplementary Figure 1**).

### Cognitive task

Each experimental session consisted of an encoding phase followed by a retrieval phase. Twenty-eight of the forty patients completed two sessions, whereas the remaining twelve patients completed more than three sessions. The original task was implemented in Chinese (**Supplementary Figure 1**), the native language of all participants; **Figure 1** shows an English-translated version for presentation purposes.

#### Encoding

During the encoding phase, participants viewed a series of 20 unique, silent video clips and were instructed to remember as much content as possible. Each trial began with a baseline fixation period during which a fixation cross was presented at the center of the screen, and participants were instructed to maintain central fixation throughout the task. The duration of this baseline period varied between 0.5 and 1.5 seconds (randomized, sampled from a uniform distribution). The fixation period was followed by presentation of a video clip that either contained no boundaries (continuous movie shots; for analytical purposes, a virtual boundary was defined at 3 seconds after clip onset) or a cognitive boundary (a cut to a new scene from a different movie, with one boundary per clip occurring 3 seconds after clip onset). Examples of clips with and without cognitive boundaries are shown in **Figure 1**. After each clip,participants pressed a button to indicate whether the majority of the clip took place indoors or outdoors.

#### Scene recognition

After completion of the encoding phase, memory for the video content was assessed using a scene recognition test. During this phase, participants viewed still frames extracted from the encoded clips (familiar frames) as well as frames taken from novel, previously unseen clips (novel frames). Participants were instructed to judge whether each frame was old or new (i.e., whether it had been seen during the encoding phase). To generate recognition stimuli, two frames were extracted from each encoded clip: one randomly selected from the first half and one from the second half of the clip. Half of these frames (n = 20) were retained as familiar frames, while the remaining half were replaced with novel frames (n = 20) drawn from different clips. The total number of familiar and novel frames per session, as well as the average similarity of novel frames, was counterbalanced between clips with cognitive boundaries and those with virtual boundaries.

### Cell classification

#### Boundary cells

For each recorded neuron, neural activity was aligned to cognitive boundaries or virtual boundaries during the encoding phase. Boundaries from all task sessions recorded within the same participant were pooled. Spike counts were computed within a post-boundary window (0–1 second) and a baseline window (−1 – 0 seconds) relative to boundary onset. A neuron was classified as a boundary cell if it satisfied both of the following criteria: (1) its firing rate in the post-boundary window differed significantly from baseline for cognitive boundary clips (*p* < 0.05, permutation test), and (2) its firing rate in the post-boundary window differed significantly between cognitive boundary clips and virtual boundary clips (*p* < 0.05, permutation test). Boundary cells were further subdivided into two types: type-I, in which post-boundary firing rates were higher for cognitive boundaries than for virtual boundaries, and type-II, showing the opposite pattern.

#### Memory cells

For memory cell identification, neural activity was aligned to image onset during the scene recognition phase, pooling trials across all sessions recorded within the same participant. Spike counts were computed in a post-image window (0– 1 seconds) and a baseline window (−1–0 seconds) relative to image onset. A neuron was classified as a memory cell if it met both of the following criteria: (1) its post-image spike counts differed significantly from baseline during novel image trials (*p* < 0.05,permutation test), and (2) its post-image spike counts differed significantly between novel and familiar image trials (*p* < 0.05, permutation test). Memory cells were further categorized as type-I, exhibiting higher post-image firing rates for novel images than for familiar images, or type-II, exhibiting the opposite pattern.

#### Chance-level estimation for cell classification

To estimate the proportion of neurons classified as boundary or memory cells expected by chance, we repeated the full classification procedures after randomly shuffling trial labels (cognitive versus virtual boundaries; novel versus familiar images) 1,000 times. For each iteration, we computed the proportion of neurons classified as boundary cells or memory cells relative to the total number of neurons recorded in the substantia nigra. These proportions formed empirically derived null distributions for boundary and memory cell classification. The observed proportions were considered significant if they exceeded the 95th percentile of the corresponding null distribution.

### Anatomical location comparison

Recording locations of overlap cells were visualized in Lead-DBS^39^ (DBS Tractography Atlas, Middlebrooks 2020^18^) by plotting cell coordinates over the SNc/SNr atlas in ICBM 2009b nonlinear asymmetric MNI space^17^. Because MER sites were obtained along bilateral trajectories and our goal was to compare overall spatial distributions rather than laterality, hemispheres were pooled by mirroring left-hemisphere coordinates across the midline (using absolute X, |X|), while preserving anterior-posterior (Y) and dorsal-ventral (Z) positions. To test whether type-I and type-II overlap cells differed in anatomical location, we performed one-way ANOVAs with neuron type (type-I versus type-II) as a between-group factor separately for each coordinate axis (|X|, Y, and Z).

### Statistical analyses

No statistical method was used to predetermine sample size, but our sample sizes are greater to those reported in previous publications^8–10,12^. Data collection and analysis were not performed blind to the conditions of the experiments. The experiments were not randomized. As mentioned above, 12 neurons were excluded from the analyses due to low firing rates. For comparisons between two conditions within the same group of cells, we used the paired *t*-test statistic. For comparisons between two different cell groups, we used a parametric one-way ANOVA. For statistical thresholding, permutation tests were conducted to generate a null distribution estimated from 1000 runs on data with scrambled labels, which avoids the assumption of normality when evaluating significance.

## Data Availability

Data that support the findings of this study will be deposited at DANDI Archive upon acceptance.

## Code Availability

Codes that support the findings of this study will be deposited at GitHub upon acceptance.

## Acknowledgements

The authors are grateful to the many patients who participated in the study, without whom this research would never have been possible, as well as numerous funding agencies for generous support, including the National Natural Science Foundation of China (82471490, L.S., 82471265, F.M.), Beijing Natural Science Foundation (JQ23038, L.S.).

