## Supplementary Tables and Figures for "Neurons in the Human Substantia Nigra Respond to Cognitive Boundaries and Predict Memory"

| Subject ID | Sex | Age | MMSE | MoCA | Recording Site | Diagnosis | UPDRS -III | UPDRS -III |
|---|---|---|---|---|---|---|---|---|
| | | | | | | | OFF | ON |
| ***Subject 01*** | F | 67 | 27(M) | 20 | L | PD | 43 | 18 |
| ***Subject 02*** | M | 49 | - | - | BiL | DT | 23 | 11 |
| ***Subject 03*** | M | 43 | 29(U) | 29 | L | PD | 34 | 17 |
| ***Subject 04*** | M | 56 | 29(U) | 28 | BiL | PD | 23 | 8 |
| ***Subject 05*** | F | 68 | 28(H) | 26 | L | PD | 46 | 32 |
| ***Subject 06*** | M | 50 | 27(U) | 25 | BiL | PD | 58 | 32 |
| ***Subject 07*** | M | 24 | 29(H) | 25 | L | DT | 51 | 41 |
| ***Subject 08*** | M | 61 | 26(H) | 21 | L | PD | 34 | 10 |
| ***Subject 09*** | M | 56 | 25(M) | 28 | BiL | PD | 57 | 14 |
| ***Subject 10*** | M | 70 | 29(M) | 26 | BiL | PD | 49 | 14 |
| ***Subject 11*** | F | 76 | 30(U) | 27 | L | PD | 40 | 18 |
| ***Subject 12*** | F | 56 | 27(H) | 28 | BiL | PD | 30 | 19 |
| ***Subject 13*** | M | 36 | 30(U) | 24 | BiL | PD | 25 | 7 |
| ***Subject 14*** | F | 66 | 25(I) | 28 | L | PD | 21 | 4 |
| ***Subject 15*** | F | 57 | 29(H) | 28 | L | PD | 57 | 14 |
| ***Subject 16*** | M | 52 | 30(M) | 29 | BiL | PD | 49 | 14 |
| ***Subject 17*** | M | 55 | 28(U) | 21 | BiL | PD | 38 | 20 |
| ***Subject 18*** | M | 47 | 29(H) | 26 | R | PD | 44 | 41 |
| ***Subject 19*** | M | 55 | 28(M) | 24 | L | DT | 39 | 16 |
| ***Subject 20*** | M | 57 | 27(M) | 23 | BiL | PD | 21 | 8 |
| ***Subject 21*** | M | 70 | 29(M) | 25 | BiL | PD | 21 | 9 |
| ***Subject 22*** | F | 62 | 27(H) | 18 | L | PD | 56 | 32 |
| ***Subject 23*** | M | 55 | - | - | BiL | PD | 63 | 38 |
| ***Subject 24*** | M | 54 | 27(H) | 24 | R | PD | 32 | 18 |
| ***Subject 25*** | F | 64 | 28(M) | 29 | R | PD | 30 | 9 |
| ***Subject 26*** | F | 45 | - | - | BiL | DT | 57 | 14 |
| ***Subject 27*** | M | 52 | 30(U) | 21 | BiL | DT | 49 | 15 |
| ***Subject 28*** | M | 57 | 25(I) | 22 | L | DT | 25 | 7 |
| ***Subject 29*** | F | 57 | 28(M) | 29 | L | PD | 21 | 4 |
| ***Subject 30*** | M | 52 | 26(U) | 23 | R | PD | 45 | 13 |
| ***Subject 31*** | M | 50 | 28(H) | 25 | L | PD | 27 | 18 |
| ***Subject 32*** | M | 67 | 28(I) | 27 | BiL | PD | 43 | 37 |
| ***Subject 33*** | F | 66 | 30(M) | 24 | BiL | PD | 18 | 4 |
| ***Subject 34*** | M | 72 | 26(M) | 24 | BiL | PD | 45 | 24 |
| ***Subject 35*** | M | 59 | 28(U) | 25 | BiL | PD | 28 | 11 |
| ***Subject 36*** | M | 56 | 28(H) | 24 | L | PD | 56 | 39 |
| ***Subject 37*** | F | 52 | 29(U) | 25 | R | PD | 27 | 13 |
| ***Subject 38*** | M | 66 | 27(H) | 22 | BiL | PD | 40 | 27 |

| | | | | | | | | |
|---|---|---|---|---|---|---|---|---|
| ***Subject 39*** | M | 69 | 26(U) | 24 | BiL | PD | 34 | 8 |
| ***Subject 40*** | M | 58 | 29(H) | 20 | BiL | PD | 39 | 28 |

**Supplementary Table 1**. Participants' demographics. The study included 40 participants with PD (26 males, 14 females) and 6 participants with DT (5 males, 1 female) who underwent DBS targeting the subthalamic nucleus. The mean age of the cohort was 57.10 years (range: 24–76 years). Cognitive assessments revealed a mean Mini-Mental State Examination (MMSE) score of 27.87 (range: 25–30) and a mean Montreal Cognitive Assessment (MoCA) score of 24.78 (range: 18–29), indicating mild to no cognitive impairment in most participants. Electrophysiological recordings were performed unilaterally in 19 patients (left: 14; right: 5) and bilaterally in the remaining 21 patients. Preoperatively, the motor symptoms, as assessed by the Unified Parkinson's Disease Rating Scale Part III (UPDRS-III), showed significant improvement post-medication (Med-ON), with scores decreasing from a mean of 38.5 (range: 18–63) in the Med-OFF state to 18.15 (range: 4–41) in the Med-ON state.

**Abbreviations:** DBS, deep brain stimulation; F, female; M, male; MMSE, Mini-Mental State Examination; I, illiteracy; H, high school; M, middle school; U, university; MoCA, Montreal Cognitive Assessment; PD, Parkinson's disease; DT, dystonia; STN, subthalamic nucleus; UPDRS, Unified Parkinson's Disease Rating Scale.

| Subject ID | MNI Coordinates (mm) | | | | | | | | |
|---|---|---|---|---|---|---|---|---|---|
| | Left Hemisphere | | | | | | | | |
| | Coordinates | | | Neuron Type | | | | Overlap | |
| | x | y | z | B-I | B-II | M-I | M-II | I | II |
| ***Subject 01*** | -9.597 | -17.1532 | -13.9 | 1 | | 1 | | 1 | |
| ***Subject 02*** | -9.5842 | -15.434 | -15.8083 | | | | | | |
| ***Subject 03*** | -10.8275 | -17.8639 | -13.3209 | | | | | | |
| ***Subject 04*** | -11.7042 | -15.41 | -12.5724 | | 1 | | 1 | | 1 |
| ***Subject 05*** | -10.7914 | -15.6888 | -13.7779 | | 1 | | 1 | | 1 |
| ***Subject 06*** | -10.7646 | -15.4584 | -14.0246 | | | | 1 | | |
| ***Subject 07*** | -10.913 | -17.5503 | -14.9852 | | | | | | |
| ***Subject 08*** | -9.9922 | -16.4243 | -15.575 | | | | | | |
| ***Subject 09*** | -8.6351 | -19.1061 | -15.5642 | 1 | | 1 | | 1 | |
| ***Subject 10*** | -9.2086 | -15.3217 | -15.1055 | 1 | | 1 | | | |
| ***Subject 11*** | -10.6742 | -18.1595 | -15.5795 | | | | | | |
| ***Subject 12*** | -10.6744 | -20.3428 | -15.7625 | | | | | | |
| ***Subject 13*** | -8.9316 | -17.5677 | -14.1734 | 1 | | 1 | | 1 | |
| ***Subject 14*** | -8.7547 | -19.2924 | -17.5751 | 1 | | 1 | | 1 | |
| ***Subject 15*** | -9.3627 | -18.2886 | -16.4977 | | | 1 | | | |
| ***Subject 16*** | -10.1213 | -19.2512 | -15.0856 | | | | | | |
| ***Subject 17*** | -9.5814 | -19.0345 | -14.5489 | 1 | | 1 | | 1 | |
| ***Subject 18*** | -9.95481 | -18.8488 | -14.3594 | | | 1 | | | |
| ***Subject 19*** | -10.0481 | -18.436 | -14.0368 | | | | | | |
| ***Subject 20*** | -10.5613 | -18.7936 | -14.6799 | | | | | | |
| ***Subject 21*** | -9.4778 | -19.6654 | -15.423 | 1 | | 1 | | 1 | |
| ***Subject 22*** | -11.5785 | -19.671 | -14.5277 | | 1 | | 1 | | 1 |
| ***Subject 23*** | -10.866 | -16.5774 | -14.9532 | | 1 | | 1 | | 1 |
| ***Subject 24*** | -9.2708 | -17.4207 | -12.668 | | | 1 | | | |
| ***Subject 25*** | -8.6525 | -19.3341 | -15.2254 | 1 | | 1 | | 1 | |
| ***Subject 26*** | -10.2131 | -18.2844 | -13.1952 | | | | | | |
| ***Subject 27*** | -9.373 | -16.5787 | -15.714 | | | | | | |
| ***Subject 28*** | -9.9874 | -18.7175 | -14.6579 | | | 1 | | | |
| ***Subject 29*** | -8.8049 | -17.4229 | -14.0908 | 1 | | 1 | | 1 | |
| ***Subject 30*** | -10.2101 | -15.4971 | -14.5277 | | 1 | | 1 | | 1 |
| ***Subject 31*** | -9.99846 | -17.0022 | -14.4894 | | | | | | |
| ***Subject 32*** | -9.3093 | -17.5521 | -14.34 | 1 | | 1 | | 1 | |
| ***Subject 33*** | -10.3306 | -18.5233 | -13.5283 | 1 | | 1 | | 1 | |
| ***Subject 34*** | -9.9247 | -18.4719 | -16.0022 | | | | | | |
| ***Subject 35*** | -9.5892 | -16.5143 | -15.7344 | | | | | | |
| ***Subject 36*** | -10.6923 | -18.2371 | -13.8528 | | | | | | |
| ***Subject 37*** | -8.2074 | -17.8552 | -13.9825 | 1 | | 1 | | | |
| ***Subject 38*** | -10.4162 | -18.2929 | -15.5251 | | | | | | |
| ***Subject 39*** | -10.3147 | -17.6341 | -14.0379 | | 1 | | 1 | | |
| ***Subject 40*** | -10.7688 | -17.4794 | -12.8173 | | | | | | |

**Supplementary Table 2**. Montreal Neurological Institute (MNI) coordinates and putative recording regions for left hemisphere substantia nigra recording during surgery.

| Subject ID | MNI Coordinates (mm) | | | | | | | | |
|---|---|---|---|---|---|---|---|---|---|
| | Right Hemisphere | | | | | | | | |
| | Coordinates | | | Neuron Type | | | | Overlap | |
| | x | y | z | B-I | B-II | M-I | M-II | I | II |
| ***Subject 01*** | 10.9883 | -23.6488 | -14.5277 | 1 | | 1 | | | |
| ***Subject 02*** | 10.376 | -16.0503 | -15.236 | | 1 | | 1 | | |
| ***Subject 03*** | 10.971 | -18.1454 | -13.9847 | | | | | | |
| ***Subject 04*** | 10.971 | -16.8185 | -14.5277 | | | | 1 | | |
| ***Subject 05*** | 10.56456 | -16.1167 | -13.0868 | | 1 | | 1 | | |
| ***Subject 06*** | 9.91161 | -16.5721 | -13.7344 | | | | | | |
| ***Subject 07*** | 10.52064 | -16.2171 | -14.0742 | | | | | | |
| ***Subject 08*** | 9.54134 | -15.9274 | -16.5222 | 1 | | 1 | | | |
| ***Subject 09*** | 10.3703 | -18.5148 | -15.0116 | | | | | | |
| ***Subject 10*** | 11.31021 | -15.5837 | -12.6784 | | 1 | | 1 | | 1 |
| ***Subject 11*** | 11.48166 | -12.6816 | -12.6784 | | 1 | | 1 | | 1 |
| ***Subject 12*** | 9.9819 | -19.5864 | -15.7554 | | | 1 | | | |
| ***Subject 13*** | 10.8398 | -18.0895 | -15.0373 | | | | | | |
| ***Subject 14*** | 8.9223 | -18.535 | -16.1875 | 1 | | 1 | | 1 | |
| ***Subject 15*** | 10.8899 | -17.6945 | -13.6596 | | | | | | |
| ***Subject 16*** | 9.4761 | -19.7719 | -15.0839 | 1 | | 1 | | 1 | |
| ***Subject 17*** | 10.1182 | -17.4182 | -13.7892 | | | | | | |
| ***Subject 18*** | 10.91583 | -18.7652 | -13.5778 | | | | | | |
| ***Subject 19*** | 9.7547 | -18.3706 | -15.0101 | 1 | | 1 | | 1 | |
| ***Subject 20*** | 10.72413 | -17.9196 | -12.7763 | | | | | | |
| ***Subject 21*** | 10.3758 | -19.1013 | -16.2558 | | | | | | |
| ***Subject 22*** | 10.6291 | -15.4302 | -13.6902 | | | | 1 | | |
| ***Subject 23*** | 10.6291 | -15.0123 | -12.639 | | 1 | | 1 | | 1 |
| ***Subject 24*** | 10.4096 | -18.8945 | -13.0751 | | | | | | |
| ***Subject 25*** | 9.66482 | -19.7557 | -16.9527 | 1 | | 1 | | 1 | |
| ***Subject 26*** | 10.54629 | -18.7838 | -13.6797 | | | | | | |
| ***Subject 27*** | 9.657 | -18.54 | -16.19 | | | | | | |
| ***Subject 28*** | 10.23147 | -17.1817 | -13.099 | | | | | | |
| ***Subject 29*** | 11.36349 | -14.8062 | -13.1829 | | 1 | | 1 | | 1 |
| ***Subject 30*** | 10.28502 | -18.045 | -13.8694 | 1 | | 1 | | | |
| ***Subject 31*** | 9.7685 | -17.9902 | -15.7509 | 1 | | 1 | | 1 | |
| ***Subject 32*** | 10.97163 | -19.5859 | -15.1734 | | | | | | |
| ***Subject 33*** | 10.81719 | -16.9117 | -13.4119 | | | | 1 | | |
| ***Subject 34*** | 9.9387 | -18.7944 | -15.1214 | 1 | | 1 | | | |
| ***Subject 35*** | 10.71468 | -15.9516 | -12.6815 | | 1 | | 1 | | 1 |
| ***Subject 36*** | 10.33902 | -17.8331 | -14.5416 | | | | | | |
| ***Subject 37*** | 9.0505 | -17.2127 | -14.1133 | 1 | | 1 | | 1 | |
| ***Subject 38*** | 10.26702 | -17.0649 | -14.143 | | | | | | |

| | | | | | |
|---|---|---|---|---|---|
| ***Subject 39*** | 10.3038 | -16.9244 | -15.2738 | 1 | 1 |
| ***Subject 40*** | 10.0566 | -19.1722 | -12.7407 | | |

**Supplementary Table 3**. Montreal Neurological Institute (MNI) coordinates and putative recording regions for right hemisphere substantia nigra recording during surgery.

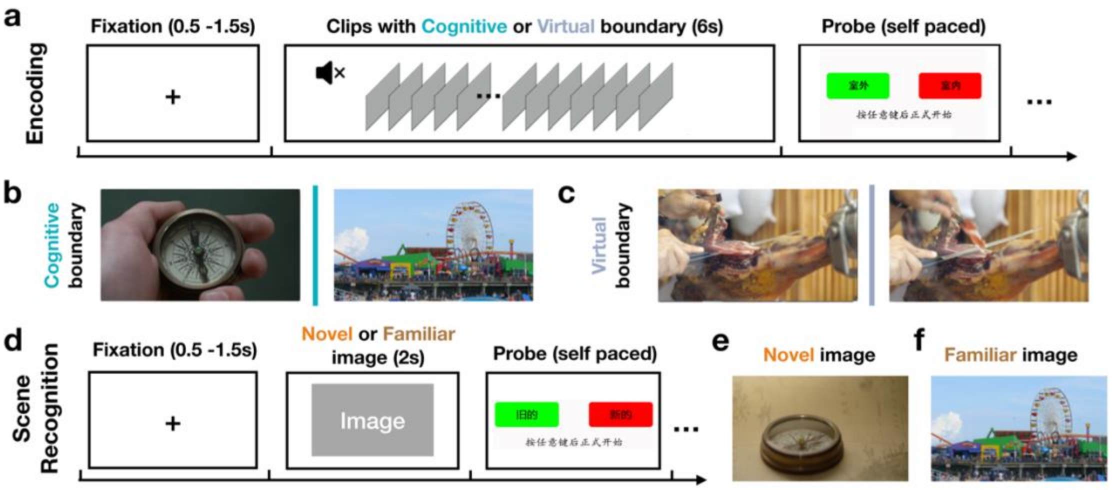

**Supplementary Figure 1 | Schematic of task presentation in Chinese. (a-c)** Encoding stage. Participants viewed a series of silent video clips and, after each clip, indicated whether most of the clip took place indoors or outdoors. Clips either contained a cut between scenes from different movies (cognitive boundary clips) or contained no cut (virtual boundary clips). For virtual boundary clips, a "virtual boundary" time point was defined at 3 s after clip onset and used as the alignment point in analyses comparing virtual and cognitive boundaries. **(b, c)** Example frames spanning a cognitive boundary (cut) and a virtual boundary (no cut) in representative clips. **(d-f)** Scene recognition stage. Participants viewed a static image and reported whether it was "old" (previously shown during encoding) or "new." **(e, f)** Example novel and familiar images. All task instructions and response prompts were presented in Chinese.

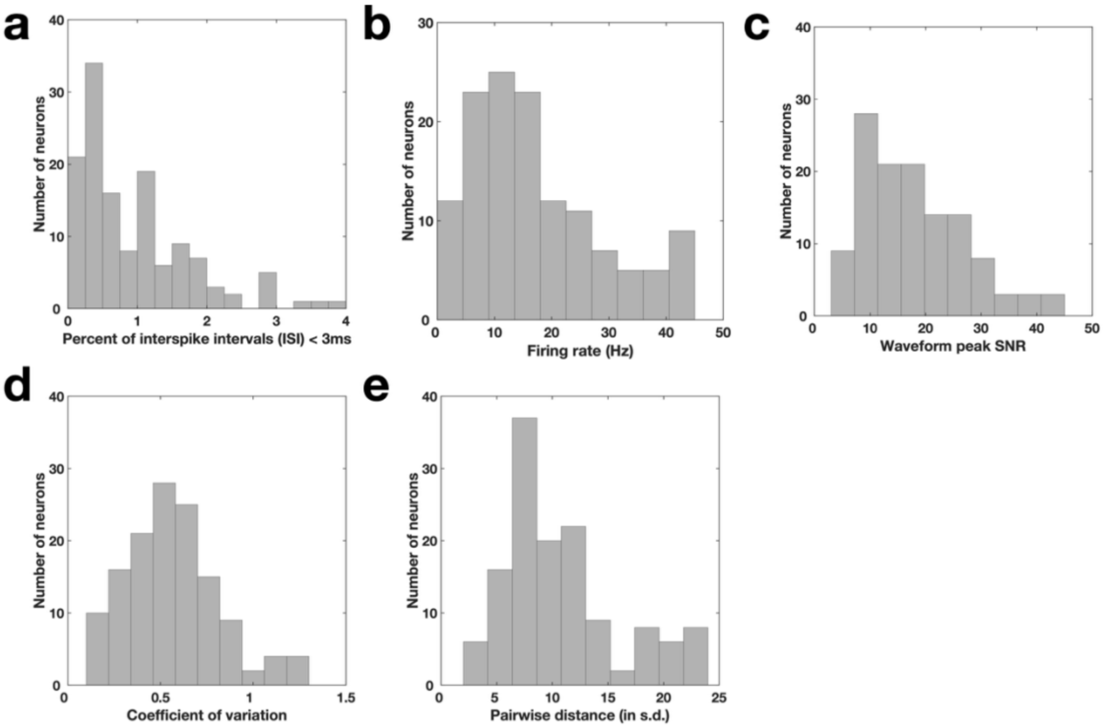

**Supplementary Figure 2 | Spike sorting quality metrics for all identified putative single cells with firing rates bigger than 0.5 Hz. (a)** Proportion of inter-spike intervals (ISI) that are shorter than 3 milliseconds. **(b)** Average firing rate within the entire recording session for all identified putative single cells. **(c)** Waveform peak signal-to-noise ratio (SNR), which is the ratio between the peak amplitude of the mean waveform and the s.d. of the noise of each identified putative single cell (8.00 ± 4,73, mean ± s.d.). **(d)** Coefficient of variation (CV2) in the ISI for each identified putative single cell (0.74 ± 0.24, mean ± s.d.). **(e)** Pairwise isolation distance between putative single cells identified from the same wire.

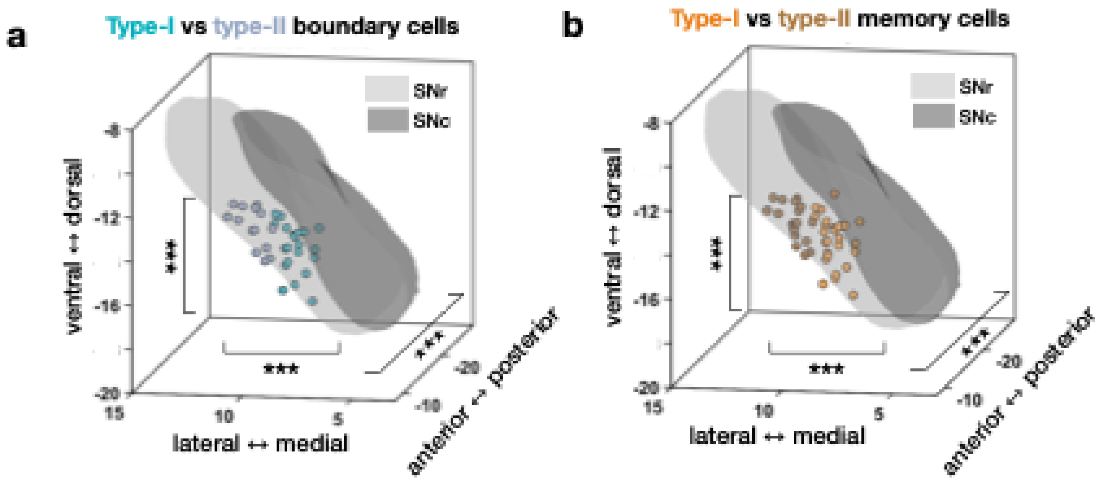

**Supplementary Figure 3 | Anatomical distributions of boundary and memory cells.** Microelectrode recording locations are shown in MNI space, with substantia nigra pars compacta (SNc) plotted in dark gray and substantia nigra pars reticulata (SNr) in light gray. **(a)** Recording sites of type-I (blue) and type-II (gray) boundary cells. **(b)** Recording sites of type-I (orange) and type-II (brown) memory cells. To pool hemispheres, left-hemisphere sites are mirrored to the right by taking the absolute value of the mediolateral (X) coordinate. Group differences along each MNI axis were assessed using one-way ANOVAs (***$p < 0.001$).
